# The cytoplasmic C-terminal domain of the MmpL11 lipid transporter is required for interaction with its co-cistronic partner MSMEG_0240 in *Mycobacterium smegmatis*

**DOI:** 10.64898/2026.01.19.699910

**Authors:** Sophie Lecher, Neetika Jaisinghani, Mary Previti, Anne-Sophie Lacoste, Jean-Michel Saliou, Dheeraj Ramchandani, Jessica C. Seeliger, Romain Veyron-Churlet

## Abstract

MmpL proteins play an important role in the various mechanisms associated with mycobacterial virulence. Identification of interacting protein partners by classical *in vitro* methods is hampered by the low solubility of these integral membrane proteins. In this study, we used two independent biotin proximity labelling assays (APEX2 and BioID) to define the proxisome of MmpL11 in *Mycobacterium smegmatis*. Crucially, these techniques are performed directly in the organism of interest, allowing the detection of potentially transient or weak interactions in multiprotein complexes and preserving the subcellular structures and the presence of cofactors or post-translational modifications that can also impact protein-protein interactions. BioID leads to the biotinylation of lysine residues, whereas APEX2 leads to the biotinylation of mainly tyrosine residues; they have also been shown to have different effective labelling radii. On one hand, an interaction was detected between the cytoplasmic C-terminal domain of MmpL11 and MSMEG_0240, a protein of unknown function, using BioID. This interaction was confirmed using both MmpL11 and MSMEG_0240 as fusions with BirA and was further supported by AlphaFold3 prediction. On the other hand, APEX2 did not detect proximity between MmpL11 and MSMEG_0240, probably due to the absence of accessible tyrosines. However, both approaches identified MSMEG_0940 as an additional interactant with MmpL11 that also depends on its C-terminal domain. Overall, this study demonstrates the utility of APEX2 and BioID as complementary tools for defining the proxisome of mycobacterial proteins.

## Introduction

The Mycobacterial membrane protein large (MmpL) proteins play critical roles in the physiology and pathogenicity of *Mycobacterium tuberculosis* (*Mtb*). For this reason, MmpL proteins have been extensively studied ^1^ ^2^ ^3^. Based on their structure, MmpL proteins have been classified into two distinct clusters ^4^. Cluster II, which includes MmpL3, MmpL11 and MmpL13, differs from cluster I by the presence of a C-terminal cytoplasmic domain that is thought to be involved in protein-protein interactions (PPIs). The C-terminal cytoplasmic domain of MmpL3 has recently been shown to interact, directly or indirectly, with Wag31 and PlrA ^5^. However, the role of the C-terminal cytoplasmic domain of MmpL11 and MmpL13 remains to be elucidated. Interestingly, MmpL11 is involved in several mechanisms related to mycobacterial virulence: cell wall biogenesis, heme transport, stress adaptation, lipid homeostasis and biofilm formation ^6^ ^7^ ^8^ ^9^ ^10^ ^11^. Accordingly, an *Mtb* mutant deleted for *mmpL11* is attenuated in a mouse model of infection, and MmpL11 is important for mycobacterial survival during later stages of chronic infection ^12^ ^13^. Thus, defining the interactome of the MmpL11 cytoplasmic domain may help to understand the role of MmpL11 in mycobacterial physiology.

New approaches to study PPIs, such as proximity-based biotinylation identification (BioID and its variants, e.g. TurboID) and APEX2, have recently been adapted to mycobacteria ^14^ ^15^ ^16^ ^17^ ^18^ ^19^. These approaches have several advantages: as they are performed directly in the organism of interest, they preserve the subcellular organisation, as well as the presence of potential post-translational modifications and cofactors. They allow the study of multiprotein complexes and can be applied to membrane-associated proteins. APEX2 and BioID differ in their application and their ability to biotinylate protein partners. While BioID is more dedicated to the characterisation of the “long history” of interactions for the protein of interest, APEX2 is more adapted to a “snapshot” of interactions, as it is performed on a shorter time scale ^20^. These approaches may be particularly useful for defining the interactome or proxisome of a given protein and for assigning a function to uncharacterised proteins in mycobacteria. For example, BioID identified an interconnexion between cholesterol catabolism and branched-chain amino acid degradation in *Mycobacterium smegmatis* (*Msm*) ^17^. APEX2 protein tagging in *Mtb* helped in localizing proteins to the cell wall or the cytoplasm, establishing the proteomes for these two subcellular compartments ^15^. In addition, APEX2 expressed as a fusion to the triacylglyceride synthase Tgs1 allowed the identification of 228 *Mycobacterium abscessus* proteins potentially involved in the intrabacterial lipid inclusion biosynthesis ^19^. Most recently, Fay *et al*. developed a two-part system based on the ALFA tag to target APEX2 to RNA polymerase RpoB and the mycolic acid synthase Pks13 and thereby identify their proxisomes ^18^. They also demonstrated successful targeting of APEX2 to the C-terminus of MmpL3, but did not identify resulting fusion-dependent labelled proteins.

In this study, a truncated version of MmpL11 (hereafter referred to as Nter), comprising the 12 transmembrane domains and lacking the cytoplasmic domain, or full-length MmpL11 were fused either to the modified biotin ligase BirA(R118G) (hereafter referred to as BirA) or to the engineered ascorbate peroxidase APEX2. Using a BioID approach in *Msm*, we defined the MmpL11 proxisome and detected an interaction between full-length MmpL11 and MSMEG_0240, a previously uncharacterised protein. We also determined that the presence of the MmpL11 cytoplasmic C-terminus was required to maintain the interaction. This interaction was confirmed by applying the BioID approach to MSMEG_0240 and further supported by AlphaFold3 prediction. This approach confirms that the cytoplasmic C-terminal domain of MmpL11 mediates PPIs. However, APEX2 did not detect an interaction between MmpL11 and MSMEG_0240, probably due to the inaccessibility of tyrosines targeted by the labelling chemistry. While most hits between the two labelling methods did not overlap, a combined analysis identified the integral membrane protein MSMEG_0940 as novel interactor based on the dependence of labelling on the MmpL11 C-terminal domain. This study has shed light on the MmpL11 interactome, the role of the C-terminal domain in PPIs, and the specific associations between MmpL11, MSMEG_0240 and MSMEG_0940, with potential implications for mycobacterial virulence and drug development.

## Results

### Production of MmpL11-BirA fusion proteins in Msm mc²155

Since BirA requires ATP to function, any PPIs involving MmpL11 can only be detected on the cytoplasmic side of the membrane in *Msm* mc²155. Therefore, BirA was fused to the N-terminus and C-terminus, both located in the cytoplasmic compartment, of truncated (Nter) or full-length MmpL11 (Figure 1A). In parallel, BirA was fused to the N-terminus of the cytoplasmic domain of MmpL11. When BirA was fused to the N-terminus of the truncated or full-length MmpL11, no transformants were obtained (data not shown), probably reflecting the toxicity of the corresponding fusion proteins in *Msm* (see Supplemental Table S1 for further details on constructs used in this study). Western blot analysis using anti-HA antibodies (when BirA was fused to the C-terminal end of truncated and full-length MmpL11) confirmed that the fusion proteins were produced in *Msm* (Figure 1B). However, the fusion proteins migrate well above their expected size (110 and 140 kDa, respectively), suggesting that MmpL11 may still be associated with mycobacterial lipids. Western blot analysis using anti-Myc (when BirA was fused to the N-terminal of the cytoplasmic domain of MmpL11) did not confirm the production of the corresponding fusion protein (data not shown), and the corresponding construct was thus not pursued further. At 16 and 40 hours, the growth of *Msm* was not affected by the production of recombinant MmpL11 or the fusion proteins with BirA, as the OD was similar for all strains of *Msm* (Figure 1C). Interestingly, treatment with the detergent dodecyl maltoside (DDM), which dissociates lipids from proteins, restored the migration of fusion proteins at their expected sizes (110 kDa for truncated MmpL11 and 140 kDa for full-length MmpL11) (Figure 1D). A greater number of additional biotinylated proteins were detected in *Msm* producing the MmpL11-fused proteins compared to the controls (empty pMV361 vector, BirA alone, or truncated or full-length MmpL11 alone), suggesting specific biotinylation of proteins closely associated with MmpL11 (Figure 1E).

**Figure 1.**
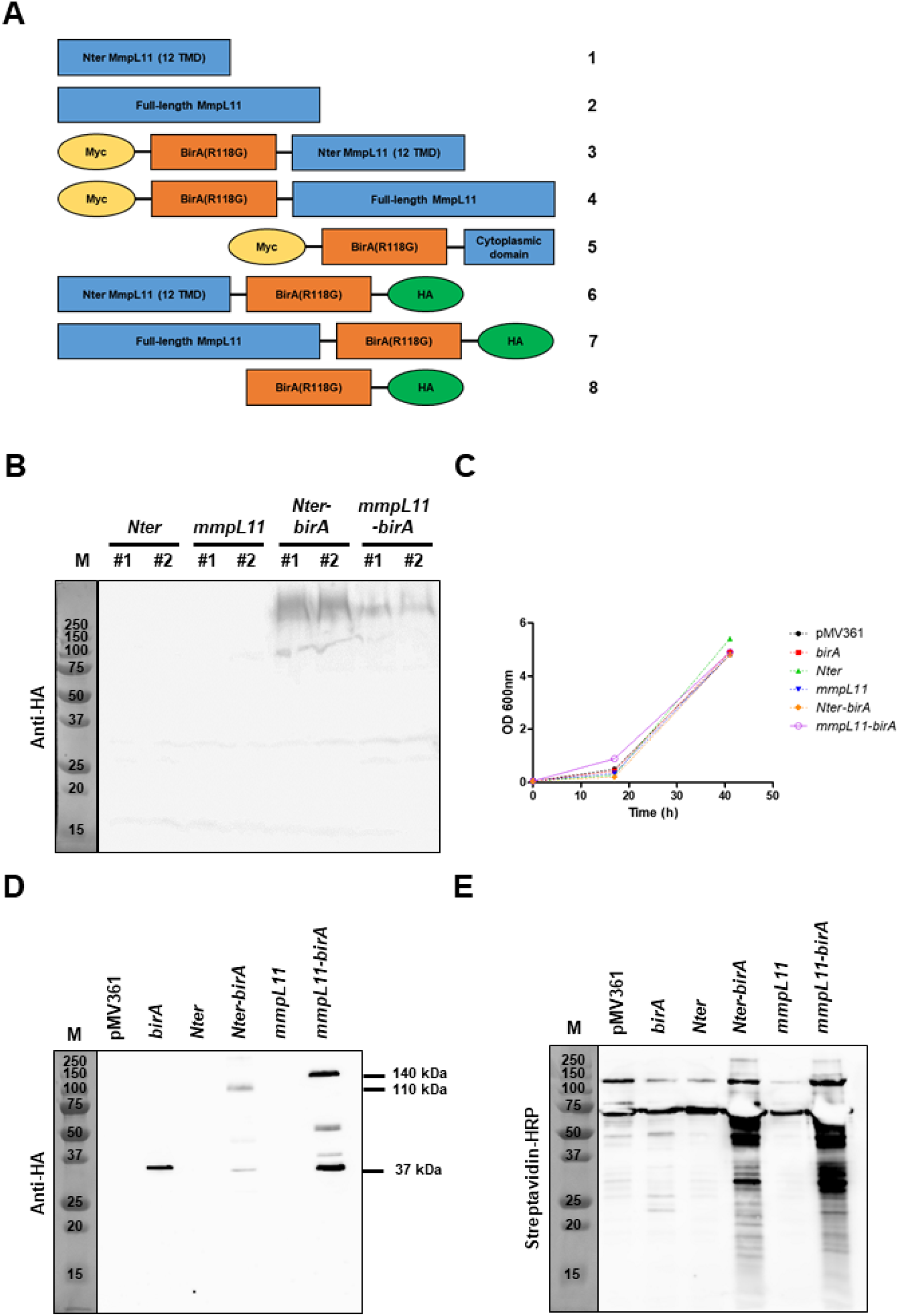
Production of fusion proteins with BirA and truncated (Nter) or full-length MmpL11 in *Msm* mc²155. **(A)** Schematic representation of the different constructs that were generated in this study. Due to toxicity and/or production issues, only constructs 1, 2, 6, 7 and 8 were further used for BioID analyses. **(B)** Anti-HA Western blot analysis of total lysates from two independent clones of *Msm* mc²155 transformed with pMV361_*Nter* (truncated), pMV361_*mmpL11* (full-length), pMV361_*Nter-birA* or pMV361_*mmpL11-birA*, as indicated. **(C)** OD_600nm_ values at 16 and 40 h from cultures of *Msm* mc²155 transformed with empty pMV361 (in black), pMV361_*birA* (in red), pMV361_*Nter* (in green), pMV361_*mmpL11* (in blue), pMV361_*Nter-birA* (in orange) or pMV361_*mmpL11-birA* (in purple). **(D)** and **(E)** Western blot analyses using anti-HA antibodies or HRP-conjugated streptavidin of biotinylated proteins from total lysates of *Msm* mc²155 transformed with pMV361, pMV361_*birA*, pMV361_*Nter*, pMV361_*Nter-birA*, pMV361_*mmpL11* or pMV361_*mmpL11-birA*, after enrichment on streptavidin beads. The molecular size markers (M) with their respective sizes expressed in kDa are shown on the left. The positions of MmpL11-BirA (140 kDa), Nter-BirA (110 kDa) and BirA (37 kDa) are indicated by the short horizontal bars. Images are representative of three independent experiments.

### Identification of MmpL11 interactants by BioID

To identify proteins that interact with MmpL11, each sample of biotinylated proteins from three independent biological replicates was analysed by mass spectrometry (Supplemental Table S3). From our previous studies and as confirmed by Western blot analyses (Figure 1E), *birA* overexpression can lead to non-specific biotinylation of mycobacterial proteins. Therefore, candidate proteins, as quantified by spectral counting, were selected by comparing their enrichment in *Msm* producing the bait of interest, truncated or full-length MmpL11 fused to BirA, versus *Msm* producing BirA alone. Only proteins that showed an enrichment ratio greater than 10 in the presence of fusion proteins compared to BirA alone were selected. In addition, since recombinant expression of *mmpL11* alone could lead to a non-specific increase in the amount of biotinylated proteins, we only considered proteins for which the MmpL11-BirA/BirA ratio was at least three times higher than the MmpL11 alone/BirA ratio.

Three interactants were enriched with the truncated version of MmpL11: MmpL11 itself, a putative fatty-acid-AMP ligase (MSMEG_4731) and the membrane-associated component EccC3 (MSMEG_0617) of the ESX-3 system (Table 1). The homotypic interaction detected between MmpL11 subunits may reflect the *in vivo* multimerization of the protein or the auto-biotinylation of the truncated MmpL11-BirA fusion. Four interactants were identified with the full-length version of MmpL11: MmpL11 itself, a conserved hypothetical protein (MSMEG_0240), EccC3 (MSMEG_0617) and the desaturase DesA3 (MSMEG_1886) (Table 1). We focused on the interaction between MSMEG_0240 and MmpL11 because it had the highest ratio among the heterotypic interactants and the corresponding genes are neighbours. Furthermore, this interaction was very specific as no spectrum corresponding to MSMEG_0240 was detected in any of the controls (pMV361 empty vector, BirA and MmpL11 alone). Since this interaction was not detected with the truncated version, we concluded that the cytoplasmic domain is important for maintaining the interaction between MSMEG_0240 and MmpL11.

**Table 1.**
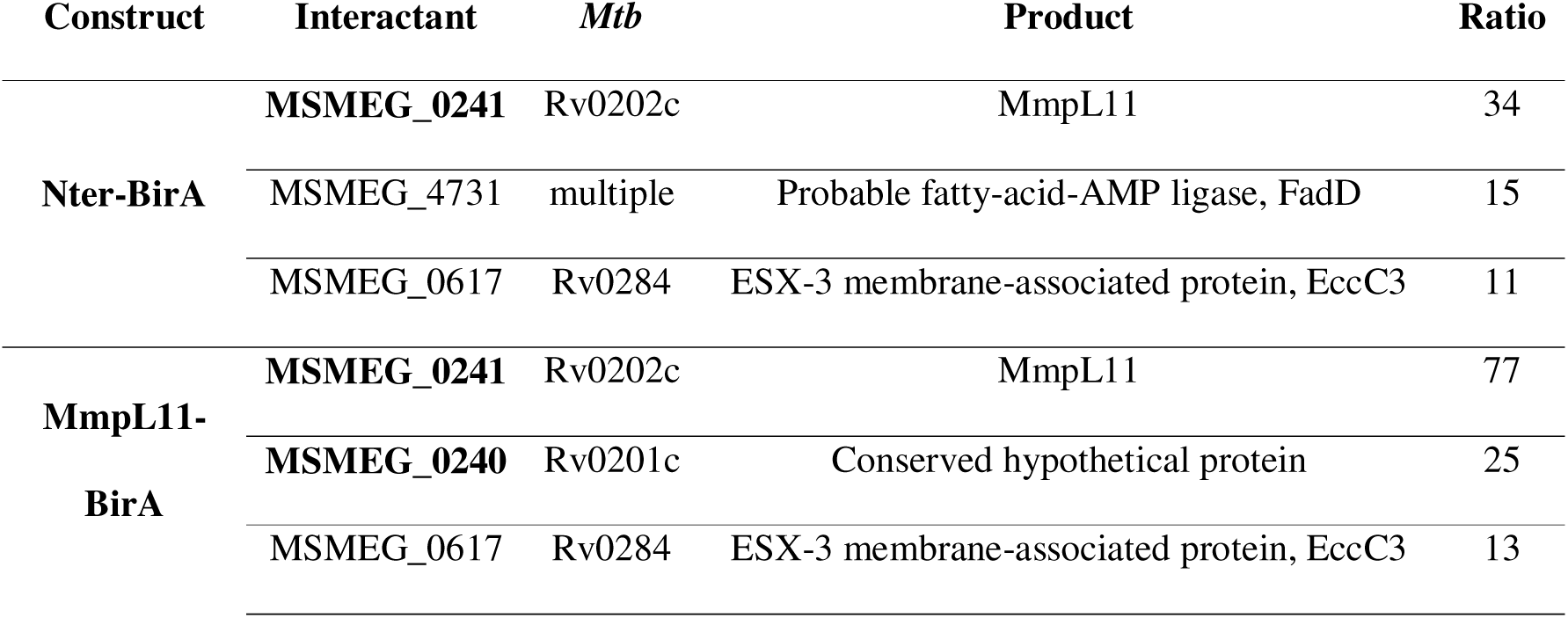
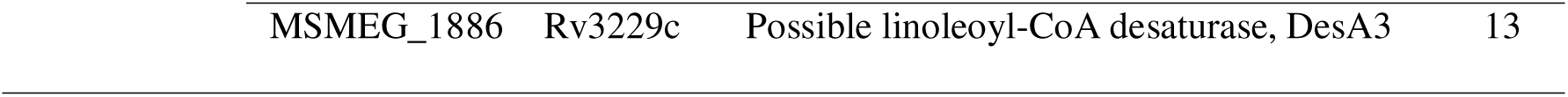
List of *Msm* proteins that interact or are closely associated with truncated (Nter) or full162 length MmpL11 fused to BirA and show a ratio > 10, when bacteria are grown in 7H9 supplemented 163 with glycerol.

### Production of MSMEG_0240 fusion proteins in Msm

To confirm further the interaction between MmpL11 and MSMEG_0240, we applied a reciprocal BioID approach using MSMEG_0240 fusion proteins with BirA (Figure 2A). Western blot analysis using anti-Myc (when BirA was fused to the N-terminal end of MSMEG_0240) or anti-HA (when BirA was fused to the C-terminal end of MSMEG_0240) antibodies confirmed that the fusion proteins were produced in *Msm* (Figure 2B). At 16 and 40 hours, the growth of *Msm* was not affected by the production of recombinant MSMEG_0240 or the fusion proteins with BirA, as the OD was similar for all strains of *Msm* (Figure 2C). Additional biotinylated proteins were detected in *Msm* producing the BirA-MSMEG_0240 fusion, but not in BirA and MSMEG_0240 alone, suggesting specific biotinylation of proteins closely associated with the BirA-MSMEG_0240 construct (Figure 2D). However, no additional biotinylated proteins were detected in the *Msm* strain producing the MSMEG_0240-BirA fusion compared to the controls.

**Figure 2.**
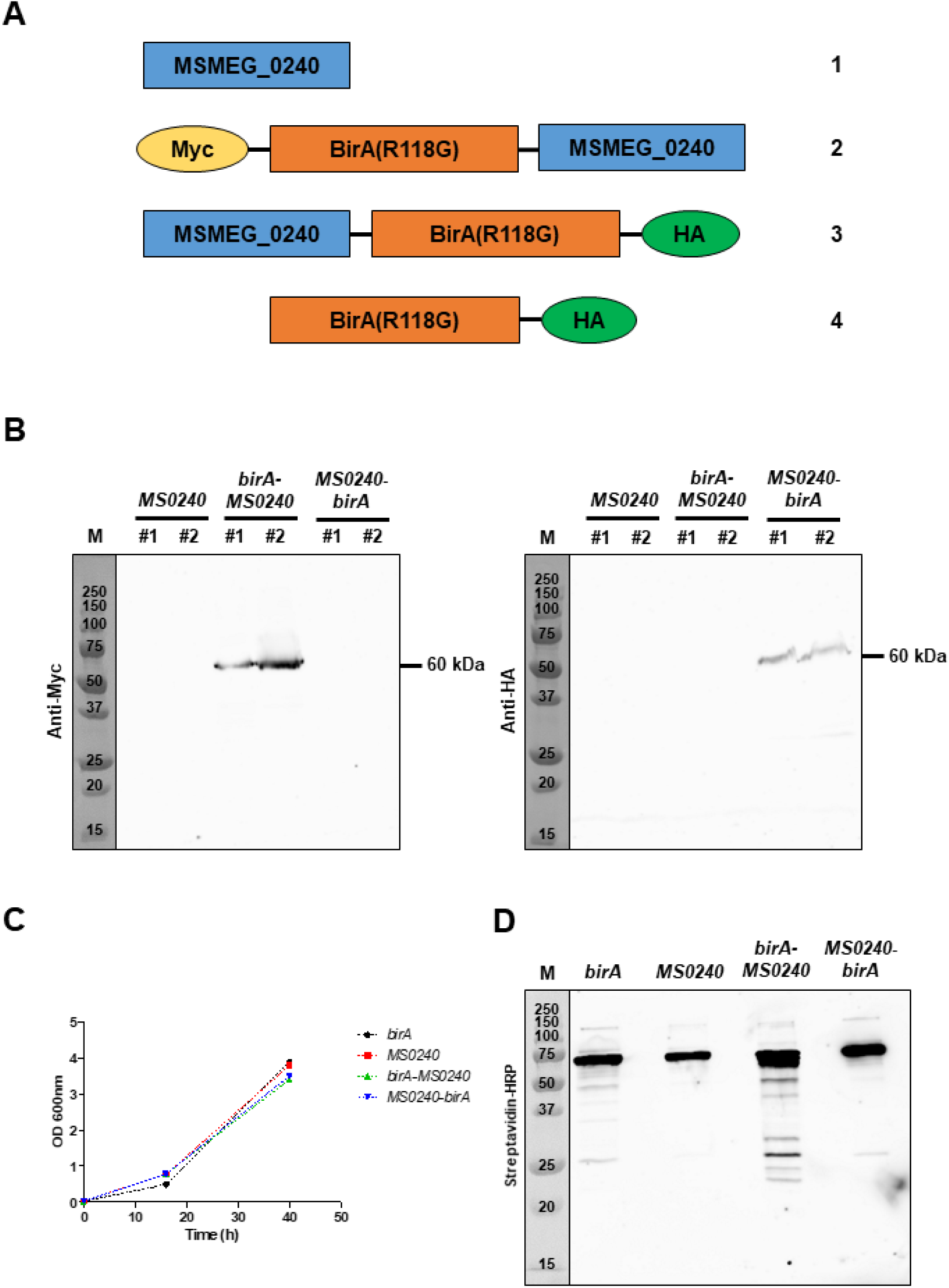
MSMEG_0240, fused to BirA, mediates biotinylation of *Msm* proteins. **(A)** Schematic representation of the different constructs generated in this study. **(B)** Western blot analysis using anti-Myc (left panel) or anti-HA (right panel) antibodies of total lysates from two independent clones of *Msm* mc²155 transformed with pMV361_*MSMEG_*0240, pMV361_*birA-MSMEG_*0240 or pMV361_*MSMEG_0240-birA*. The positions of BirA-MSMEG_0240 (60 kDa) and MSMEG_0240-BirA (60 kDa) are indicated by the short horizontal bars. **(C)** OD_600nm_ values at 16 and 40 h from cultures of *Msm* mc²155 transformed with pMV361_*birA* (in black), pMV361_*MSMEG_*0240 (in red), pMV361_*birA-MSMEG_*0240 (in green) or pMV361_*MSMEG_0240-birA* (in blue). **(D)** Western blot analysis using HRP-conjugated streptavidin of biotinylated proteins from total lysates of *Msm* mc²155 transformed with pMV361_*birA*, pMV361_*MSMEG_*0240, pMV361_*birA-MSMEG_*0240 or pMV361_*MSMEG_0240-birA*, after enrichment on streptavidin beads. The molecular size markers (M) with their respective sizes expressed in kDa are shown on the left. Images are representative of three independent experiments.

### Identification of MSMEG_0240 interactants by BioID

To identify proteins that interact with MSMEG_0240, each sample of biotinylated proteins from three independent biological replicates was analysed by mass spectrometry (Supplemental Table S4). In agreement with previous western blot analyses (Figure 2B, right panel, and Figure 2D), the MSMEG_0240-BirA fusion was poorly detected by mass spectrometry (1 or 2 spectra), which did not allow us to detect any interactant. On the other hand, the BirA-MSMEG_0240 fusion was well produced and detected (between 25 and 40 spectra per replicate) and five interactants were detected: MSMEG_0240 itself, MmpL11, MSMEG_3471, MSMEG_5468 and MSMEG_6863 (Table 2). BirA-MSMEG_0240 was able to interact with itself, suggesting *in vivo* multimerization of the MSMEG_0240 or auto-biotinylation of the fusion protein. Furthermore, the interaction between MSMEG_0240 and MmpL11 was confirmed, using either full-length MmpL11 or MSMEG_0240 as bait. Thus, the use of a BioID approach represents a novel and robust tool for establishing the proxisome of mycobacterial membrane-associated proteins, such as MmpL proteins.

**Table 2.**
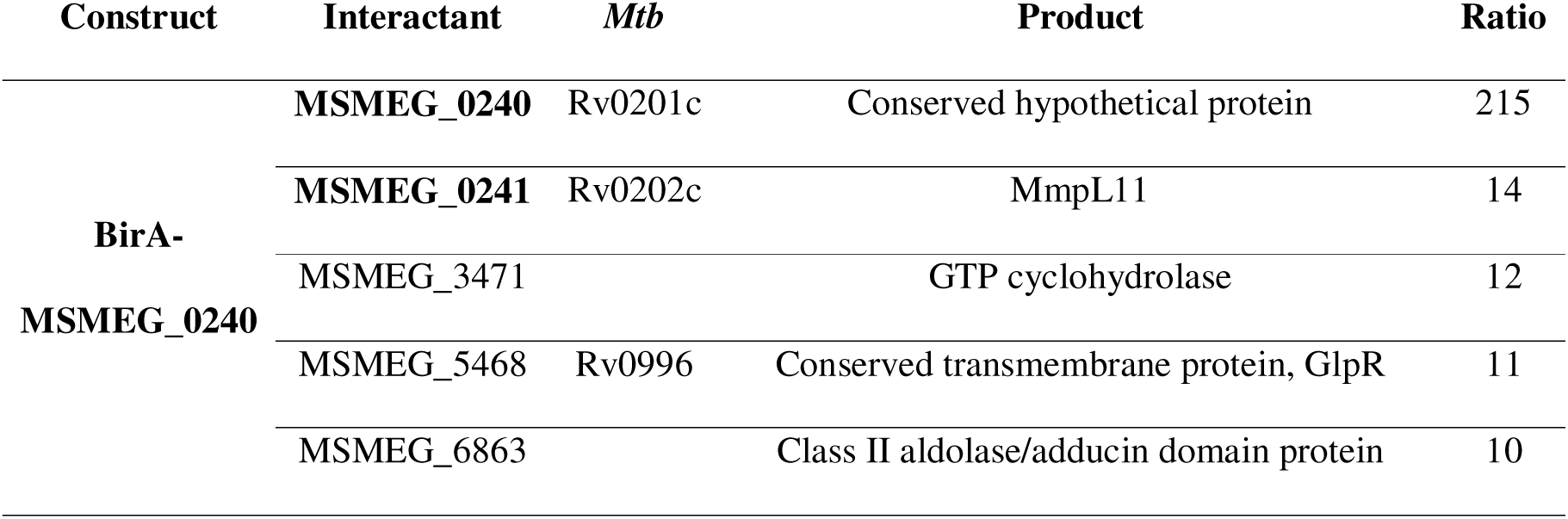
List of *Msm* proteins that interact or are closely associated with BirA-MSMEG_0240 and have a ratio > 10, when bacteria are grown in 7H9 medium supplemented with glycerol.

### AlphaFold3 supports an interaction between MmpL11 and MSMEG_0240

The gene *MSMEG_0240* is located downstream of the gene encoding for MmpL11 (*MSMEG_0241*) and they are separated by only two base pairs on the *Msm* mc²155 chromosome. The corresponding protein exhibits some homology with the transcriptional regulators of the XRE family and contains a putative helix-turn-helix motif. This genetic organisation is also present in *Mtb* H37Rv, where the orthologous gene (*rv0201c*) is located downstream of *mmpL11* (*rv0202c*), with an overlap of four base pairs. Therefore, the interaction between MSMEG_0240 and MmpL11 may support a mechanism common to the *Mycobacterium* genus. At the protein level, MSMEG_0240 contains an additional N-terminal extension, which is absent from Rv0201c and which is predicted to be largely unstructured. Full-length MmpL11 and a truncated version of MSMEG_0240 (from Asp 61 to Lys 202) were submitted to an AlphaFold3 prediction ^21^. Consistent with our results from BioID (Tables 1 and 2), an interaction was predicted between the cytoplasmic domain of MmpL11 and MSMEG_0240 (Figures 3A, 3B and 3C), with pTM and ipTM scores of 0.61 and 0.74, respectively. This model is conserved across different mycobacteria (Supplemental Figure S1). Furthermore, the putative helix-turn-helix motif of MSMEG_0240 was outside of the predicted interaction interface between MSMEG_0240 and MmpL11 (Figures 3A and 3B).

**Figure 3.**
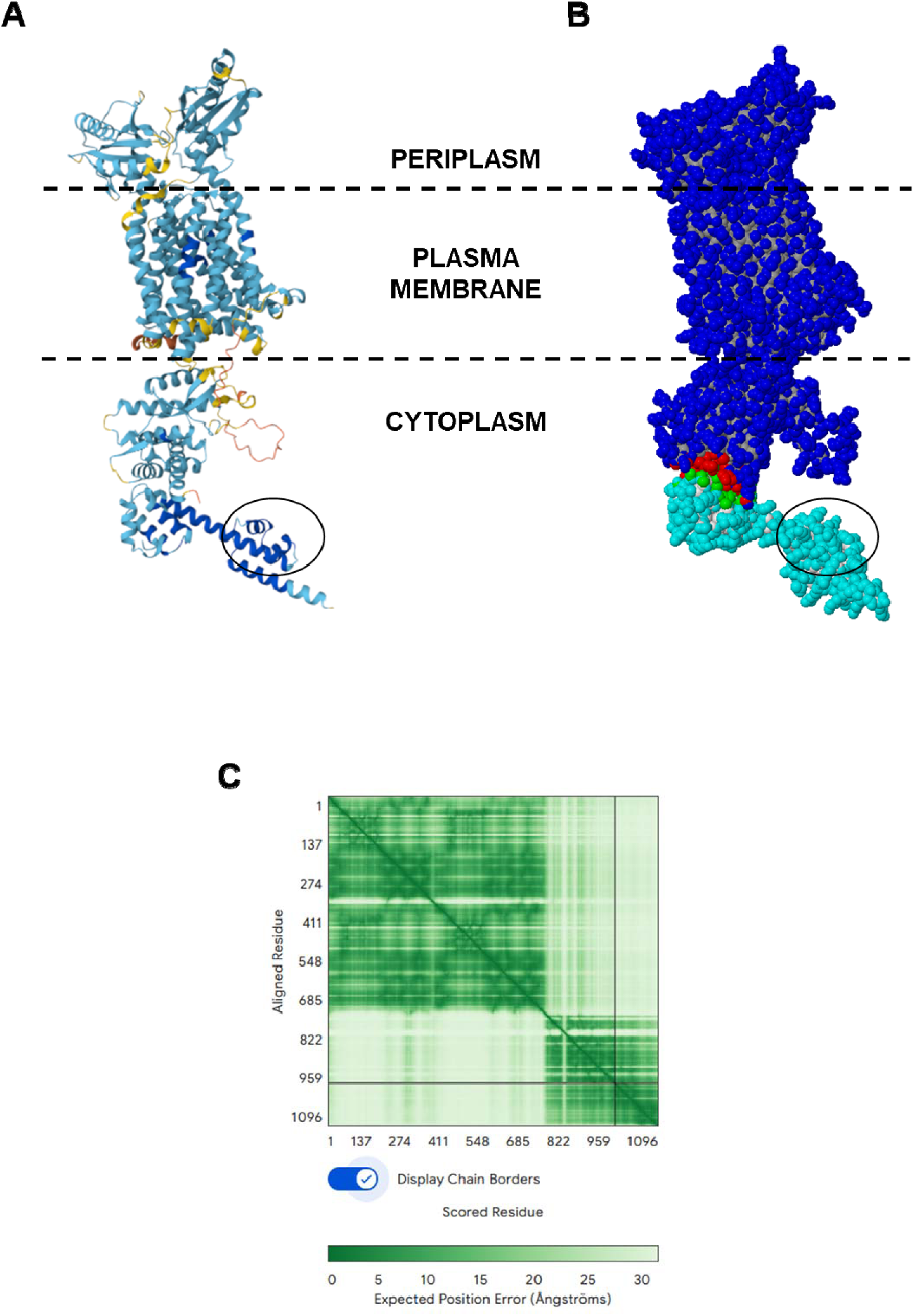
The interaction between MSMEG_0240 and the cytoplasmic domain of MmpL11 is confirmed by AlphaFold3 prediction. **(A)** AlphaFold3 prediction of the interaction between a truncated version of MSMEG_0240 (from Asp 61 to Lys 202) and the cytoplasmic domain of MmpL11^21^. The putative position of the plasma membrane is delineated by dash lines, based on the extent of the MmpL11 transmembrane helices. The structure of the proteins is labelled in dark blue (very high, pLDDT > 90), light blue (high, 90 > pLDDT > 70), yellow (low, 70 > pLDDT > 50) and orange (very low, pLDDT < 50) according to the model confidence. **(B)** Schematic representation of truncated MSMEG_0240 (light blue) and MmpL11 (dark blue) from PISA ^22^. The interacting atoms are shown in green for MSMEG_0240 and in red for MmpL11. The localisation of the putative helix-turn-helix motif of MSMEG_0240 is circled in black and is outside the predicted interaction interface. **(C)** Predicted aligned error (PAE) for the predicted interaction between MmpL11 (residues 1 to 954) and MSMEG_0240 (residues 955 to 1096).

### Identification of MmpL11 interactants by APEX2

Given the few hits detected from BirA fusions (Tables 1 and 2), we posited that the apparent proximal proteome was limited by the labelling method. We thus sought to expand the set of MmpL11 interactors using APEX2 fusions and also to provide a head-to-head comparison of the biotin ligase and peroxidase approaches. Tagging by APEX2 is expected to provide an independent measure of a proximal proteome because (1) APEX2 provides a snapshot of nearby proteins at the time of labelling since the reaction is initiated by the addition of a phenol substrate and hydrogen peroxide and proceeds for an experimentally determined time period, (2) the labelling radius of APEX2 is estimated to be different than that of BirA, and (3) APEX2 labels aromatic amino acids, understood to be primarily tyrosines, rather than lysines as for BirA.

We previously established APEX2 as a compartment-specific protein tagging method for the cytosol and periplasm of mycobacteria ^15^. Extending this earlier work, we constructed APEX2 fused to either MmpL11 or Nter, as above with BirA (Figure 4A; see also Supplemental Table S1). As we have done previously, both fusions were expressed under translational control of a theophylline-inducible riboswitch and without an additional epitope tag, as the fusions can be confirmed with a native antibody against the plant peroxidase APX2 (Figure 4B). We found that expression from an integrated single-copy plasmid did not yield obvious APEX2-dependent labelling by biotin phenol, as detected by avidin (data not shown). Expression from a multicopy (3-5 per cell) episomal plasmid, however, yielded biotinylated proteins, with distinct profiles for fusion to MmpL11 versus Nter (Figure 4C). Overall, MmpL11-APEX2 yielded more abundant and promiscuous labelling than Nter-APEX2. To check for artifacts due to overexpression of MmpL11- or Nter-APEX2 alongside the native copy of MmpL11, we also expressed both fusions in a targeted Δ*MSMEG_0241* (Δ*mmpL11*) null strain ^10^. Expression of the APEX2 fusions in either the wild-type or knockout yielded similar patterns of labelling, supporting the use of either strain to detect MmpL11-proximal proteins (Figure 4B).

**Figure 4.**
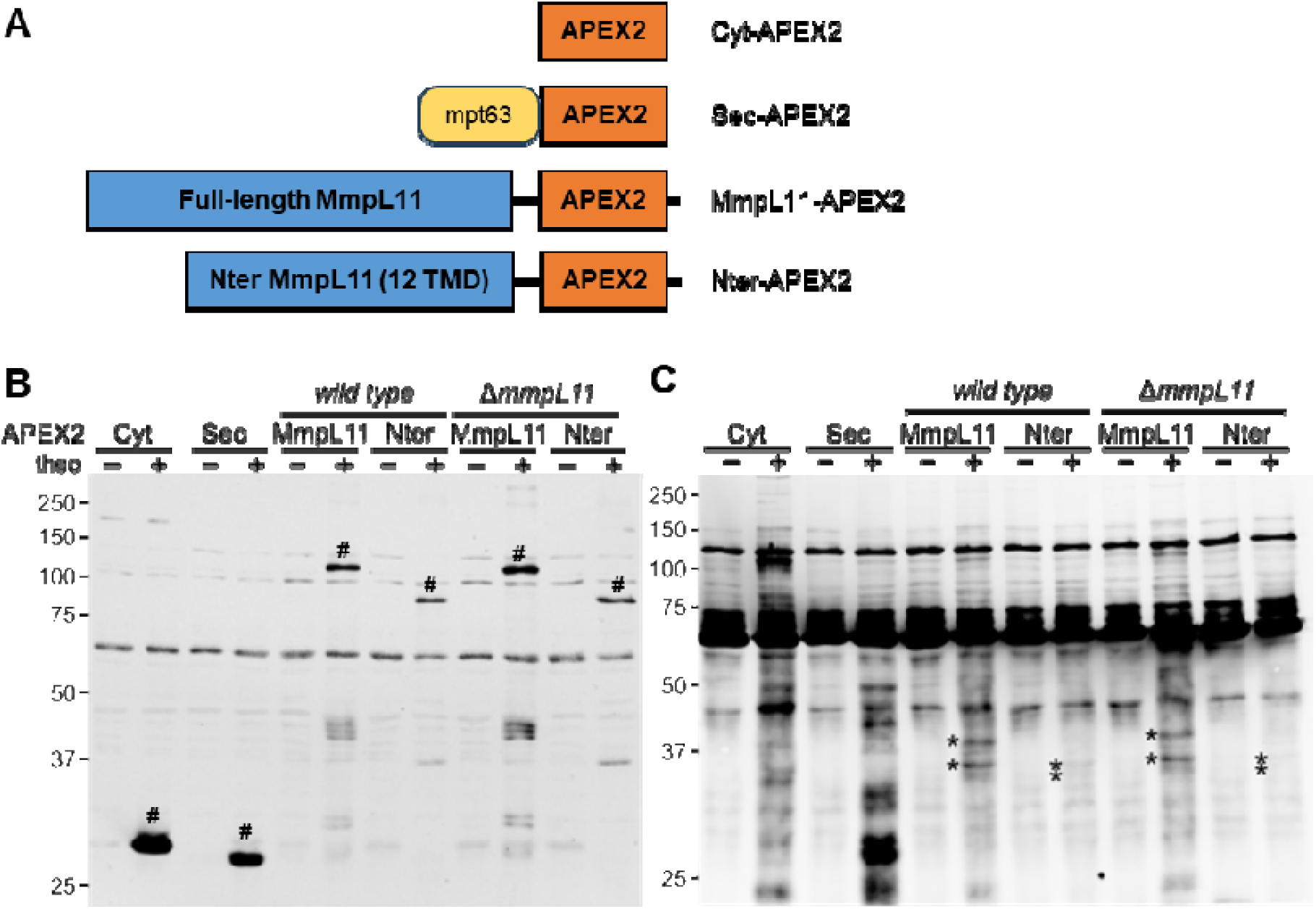
MmpL11 fused to APEX2 yields distinct protein labelling in *Msm*. *Msm* encoding APEX2 in the cytoplasm (Cyt) or cell wall (Sec), or fused to full-length MmpL11 (MmpL11) or N-terminal MmpL11 (Nter) was cultured without (−) or with (+) theophylline inducer (theo) for 6h and subjected to the labelling protocol with biotin-phenol. MmpL11 fusions were also expressed in *Msm* Δ*mmpL11* as indicated. **(A)** Schematic of APEX2 constructs used in this study. For Sec-APEX2, “*mpt63*” indicates an N-terminal Sec secretion signal, as described^15, 23^. **(B)** APEX2 expression detected by anti-APX2 immunoblot with # indicating bands with expected migration for the corresponding APEX2 construct. **(C)** APEX2-dependent labelling detected by an avidin conjugate. Asterisks indicate representative bands that are dependent on MmpL11- or Nter-, but not Cyt- or Sec-APEX2. Loads were normalized to 10 µg total protein per lane. Blots are representative of n = 3 independent experiments.

We went on to enrich labelled proteins by avidin affinity and to identify isolated proteins compared to uninduced controls by mass spectrometry. We also performed these experiments for cytosolic (Cyt) and cell wall (Sec) APEX2 to identify the corresponding subcellular proteomes, as we have done in *Mtb* ^15^. For consistency and based on the similarity of labelling between wild-type and Δ*MSMEG_0241* strains (Figure 4B), the wild type was used as the parent strain for all APEX2 constructs (Supplemental Table S5). We used SAINT analysis ^24^ to identify APEX2-dependent enriched proteins from MS/MS data, enabling comparison with analogous results in *Mtb*. To facilitate analysis, we developed a publicly available web interface that allows users to run SAINTexpress remotely rather than requiring local installation (https://www.saintexpress.org/; see Methods).

For each analysis we determined an appropriate SAINT score cutoff by comparing the numbers of hits as a function of cutoff value and found that in general, the number of hits increased sharply between SAINT score 0.7 and 0.6, suggesting non-specific detection within in this stratum. We thus used a cutoff of SAINT ≥ 0.7 or 0.65 depending on the analysis and discuss exceptions as relevant below (Supplemental Table S6). Finally, based on the initial BirA analysis above, the C-terminus of MmpL11 is critical for interaction with MSMEG_0240. We thus also applied SAINT analysis to identify detected proteins that were enriched after labelling by MmpL11- versus Nter-APEX2 fusions and thereby reveal the proteins whose proximity to MmpL11 depend the C-terminal domain (Supplemental Table S6).

While the number of proteins identified by Cyt-APEX2 was similar between *Msm* as determined here and *Mtb* as we reported previously (457 and 486, respectively), Sec-APEX2 labelling yielded only about half as many hits in *Msm* versus *Mtb* (126 and 254) (Figure 5A) ^15^. Given that the same expression constructs were used in both species, the reason for this difference is unclear. It is unlikely that there are fewer proteins in the *Msm* periplasm, especially given that *Msm* has a larger genome and encodes many more predicted solute transporters and corresponding solute binding proteins ^25^. As in *Mtb* we found that proteins identified by Sec-APEX2-dependent tagging are enriched for predicted secretion signals and transmembrane helices compared to those enriched from Cyt-APEX2 (Figures 5A and 5B), supporting labelling specifically within the cell wall ^15^. In contrast, hits dependent on MmpL11-APEX2 and Nter-APEX2 are predicted to be cytosolic or have transmembrane helices, consistent with the expected position of C-terminally fused APEX2 proximal to the cytoplasmic leaflet of the plasma membrane (Figures 3 and 5B).

**Figure 5.**
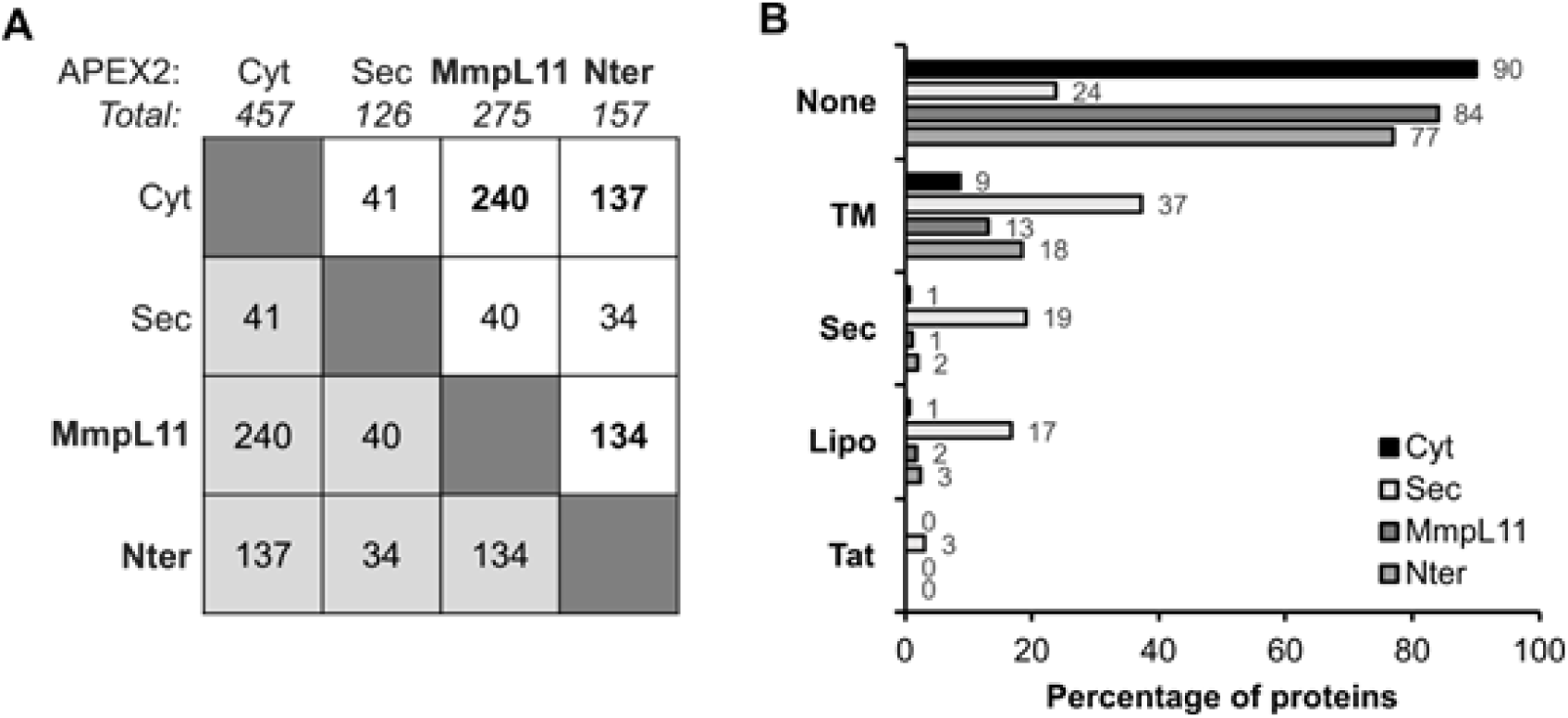
MmpL11-APEX2 fusions label a subset of cytoplasmic and transmembrane proteins. Proteins were enriched by avidin affinity from APEX2-expressing *Msm* strains labeled with biotin-phenol and identified by quantitative label-free proteomics. (A) Number of proteins identified by each APEX2 construct and their pairwise overlap. (B) Protein hits were analyzed for predicted transmembrane helices (TM), secretion signal motifs (Sec, Tat), or lipoprotein modification motif (Lipo). Proteomics analyses were derived from n = 3 independent replicates for each sample group.

Also, as in *Mtb*, APEX2-mediated labelling in *Msm* identified ESX-1 Type VII secretion system components or substrates in the cytosol by Cyt-APEX2 (EspK, EspB substrates) or periplasm by Sec-APEX2 (EccB1, MycP1 components) as consistent with the topology of the core ESX complex. However, unlike in *Mtb*, detection in *Msm* was limited to only one ESX system (ESX-1) versus three in *Mtb* (also ESX-2 and ESX-5). *Msm* does not encode ESX-2 or ESX-5 homologous systems, but does have ESX-3 and ESX-4, components of which were not detected by APEX2. In contrast, both MmpL11 fusions to BirA identified the ESX-3 core component EccC3 (Table 1), possibly underscoring the different labelling chemistry (frequency of accessible Lys vs. Tyr). While a functional underpinning for MmpL-ESX interaction is not obvious, we note that we previously detected crosslinking between another lipid transport-related protein, LprG, and components of the Sec secretion system^26^. Since mycobacteria grow from their poles, the close proximity of protein secretion and lipid transport systems may be related to synthesis of the complex multilayer cell envelope and the need to export proteins directly where they are needed. Also, detection of substrates such as EsxB in the periplasm was a major outcome in *Mtb*, but was not recapitulated in *Msm*. The lower number of proteins identified by Sec-APEX2 in *Msm* may indicate overall lower sensitivity of detection in this compartment and explain the more limited detection of ESX components. Finally, the trehalose monomycolate transporter MmpL3 (MSMEG_0250) was enriched from MmpL11-APEX2 labelling, but not by any other APEX2 construct (Supplemental Table S6). This interaction has precedent in a previous MmpL3 interactome study, in which Belardinelli *et al*. detected MmpL11 as an MmpL3 interactor by bacterial two-hybrid in *E. coli* ^27^. This same study did not detect any interactions between MmpL3 and central mycolic acid biosynthesis proteins. In contrast, proximal proteins identified by MmpL11-APEX2 (but not Nter-APEX2) included not only MmpL3, but also proteins screened by Belardinelli *et al*. (KasA/MSMEG_4327, UmaA/MSMEG_0913, FadD32/MSMEG_6393) and the additional mycolic acid modification enzymes DesA1 and DesA2 (MSMEG_5773, MSMEG_5248) (Supplemental Table S6). These results together support interactions between mycolic acid biosynthesis and transport machinery and MmpL11 that depends on its C-terminal domain.

### Comparison between BioID and APEX2 results

We further re-analyzed the MmpL11-BirA, BirA-MSMEG_0240 and MSMEG_0240-BirA datasets (Supplemental Tables S3 and S4) from above by SAINT analysis. We note that each MmpL11-APEX2 fusion yielded ∼150-250 protein hits (Supplemental Table S6), whereas by both fold-change and SAINT analysis, BirA fusions detected only 4-5 proteins (Supplemental Tables S7 and S8; note that a cutoff of 0.31 was used for BirA-MSMEG_0240 based on this SAINT score for *bona fide* interactor MmpL11/MSMEG_0241). This difference is perhaps unexpected given the shorter labelling time and lower frequency of target residues for APEX2 versus BirA. However, experimental estimates suggest that the radius of labelling by APEX2 is at least twice as large as by BirA ^28^ ^29^. Also, while the relative expression levels of the APEX2 and BirA fusions was not determined, all else being equal, higher expression is expected for APEX2 from a multi-copy compared to BirA from a single-copy vector. These factors may account for the relative promiscuity of APEX2-dependent labelling.

As was observed for labelling by BirA, MmpL11 itself was a significant hit by APEX2. However, MSMEG_0240 was not detected in any APEX2 dataset. To test APEX2 labelling of MSMEG_0240 directly, we co-expressed MmpL11-APEX2 and Nter-APEX2 with the genomic copy of MSMEG_0240 fused to a C-terminal FLAG epitope. Although the MSMEG_0240-FLAG fusion was detected in total lysates, it was not enriched by avidin after APEX2 labelling (Supplemental Figure S2). Together, the MS and affinity enrichment data suggest that the four tyrosines in MSMEG_0240 are not accessible to tagging by biotin phenoxyl radicals generated by APEX2. Based on the AlphaFold3 model generated for Figure 3, MSMEG_0240, Tyr96 and Tyr170 are involved in core interactions between helices and Tyr113 is buried in the interface with the MmpL11 C terminus, likely making these three residues inaccessible to labelling. The remaining Tyr66 is proximal to the unstructured N-terminus (residues 1-61). One hypothesis is that this unstructured region is involved in PPIs and that Tyr66 is inaccessible to labelling when MSMEG_0240 is bound. In contrast, all seven lysines of MSMEG_0240 are at the surface and thereby accessible to BirA tagging. This structural analysis provides a probable explanation for selective tagging of MSMEG_0240 by BirA versus APEX2.

For MmpL11-BirA, both fold-change and SAINT analysis yielded MmpL11 itself and MSMEG_0240 as the most significant hits, but other hits did not overlap between the two analysis methods (Tables 1 and 2; Supplemental Tables S6 and S7). However, by SAINT analysis, MSMEG_0940 is a hit from MmpL11 fused to either APEX2 or BirA, and also when comparing MmpL11-APEX2 to Nter-APEX2. Detection across these multiple methods and comparisons suggests that MSMEG_0940 is a *bona fide* interactor whose proximity depends on the presence of the MmpL11 C-terminus. MSMEG_0940 (*Mtb* homologue: Rv0497) is a transmembrane protein of unknown function with three transmembrane helices at the C-terminus; the N-terminus is largely unstructured, with little secondary structure of low confidence. In both *Msm* and *Mtb*, this protein appears to be in an operon with a predicted apurinic/apyrimidinic (AP) endonuclease (MSMEG_0941; Rv0498). AP endonucleases traditionally have roles in DNA repair by cleaving the DNA backbone at lesion sites. The possible functional connection between this predicted activity and the co-cistronic transmembrane protein MSMEG_0940 is not obvious, nor is a possible functional connection to MSMEG_0240 as a putative transcription factor and MSMEG_0241/MmpL11 as a lipid transporter. The human AP endonuclease APE1/Ref-1 has been implicated in the redox regulation of transcription factors through one of its cysteine residues (Cys65) and interaction with transcription factors via a cognate cysteine ^30,31^. MSMEG_0941 has three cysteines, including one at a similar position (Cys 68) whose position is conserved in the homologue Rv0498. However, MSMEG_0240 does not appear to be a candidate substrate in a redox mechanism, as it lacks cysteines.

## Discussion

The interactome of MmpL3 has already been determined using classical approaches such as the bacterial two-hybrid system, which provides valuable information on its function ^27^. However, two-hybrid approaches are performed in heterologous hosts (e.g. *E. coli* or yeast) and only allow the detection of pairwise interactions. The biotinylation-based proximity assays, BioID and APEX2, can be performed directly in the host of interest (here mycobacteria) and allow the study of multiprotein complexes, while preserving the subcellular organisation, the presence of cofactors and the translational modifications. In addition, these approaches allow the study under different conditions that can modify the profile of PPIs (medium composition, bacterial growth phase, temperature, etc.). Complementary to this work, we here used both MmpL11 and MSMEG_0240 as baits and showed that these two proteins interact with each other in a manner dependent on the MmpL11 C-terminal domain, further underscoring the value of BioID in dissecting the interactions and thereby the potential roles of membrane-associated or uncharacterised proteins in mycobacteria. In addition, we used MmpL11 as a system to directly compare the proteomes detected by BioID versus the peroxidase APEX2, which involves distinct labelling chemistry. Consistent with this difference, MSMEG_0240 was not detected by MmpL11-APEX2, likely due to the inaccessibility of target tyrosines. On the other hand, the detected proteome was two orders of magnitude larger than for BirA fusions. This may suggest that BirA labelling in this context is more specific, giving robust but incomplete information. On the other hand, APEX2 may be more non-specific due to a larger radius of labelling and thus less informative for inferring functional relationships between co-localized proteins. Most informative is the comparison of MmpL11 and Nter fusions, where dependence on the C-terminus increases confidence in the specificity of the identification. Overall, these comparisons provide experimentally observed differences between biotin ligase and peroxidase-mediated labelling that can inform their application to targets in mycobacteria. In particular, the use of APEX2 in a protein domain-dependent or other structure-dependent context may provide the most accurate and comprehensive picture of an interactome.

Based on the presence of a putative helix-turn-helix motif, MSMEG_0240 shows homologies with the XRE family of transcription factors. The corresponding motif is outside from the predicted interface of interaction with MmpL11 (Figures 3A and 3B), inferring that MSMEG_0240 may bind DNA while interacting with MmpL11. However, the DNA-binding ability of MSMEG_0240, with or without MmpL11, remains to be assessed. If so, the organisation of MmpL11 and MSMEG_0240 would be reminiscent of sigma/anti-sigma factors or two-component systems of mycobacteria. Transcriptomic analyses comparing the wild-type strain of *Msm* with the *MSMEG_0240* deletion mutant may uncover genes regulated by this protein. In addition, the overexpression of full-length or truncated MmpL11 in a *mmpL11* deletion mutant could reveal genes whose expression is regulated by MSMEG_0240 in an MmpL11 C-terminus-dependent manner. Moreover, the function of MmpL11 in *Mtb* has been shown to be regulated by phosphorylation on its cytoplasmic C-terminal domain ^11^. Interestingly, a phosphorylation site in *Mtb* (serine 956) is conserved in *Msm* (serine 944) and is located at the predicted interface of the interaction between MmpL11 and MSMEG_0240. Thus, phosphorylation of this residue may directly regulate the interaction between MmpL11 and MSMEG_0240.

Finally, the characterisation of the interaction between MmpL11 and Rv0201c, the MSMEG_0240 orthologue in *Mtb*, may be of therapeutic interest. A transposon library screen indicated that MmpL11 and Rv0201c are each required for intracellular survival of mycobacteria in J774 murine macrophages ^32^. Furthermore, cefoxitin is an intravenous beta-lactam used in the intensive treatment phase of human pulmonary infections caused by *Mycobacterium abscessus* and very recent work has shown that cefoxitin-resistant clones of *M. abscessus* carry a mutation in either MmpL11 or the MSMEG_0240 orthologue ^33^, suggesting that cefoxitin may disrupt the relationship between the two proteins. Alternatively, the interaction between MmpL11 and the MSMEG_0240 orthologue may be required for the efficient transport of cefoxitin, in a mechanism similar to that known for other MmpL proteins, such as MmpL5 and bedaquiline efflux in *Mtb* ^34^. In addition, another study has also shown that mutations in MmpL11 or MSMEG_0240 confer resistance to fluoroquinophenoxazine, a putative non-beta-lactam topoisomerase inhibitor in *Msm* ^35^. Thus, interfering with the complex formed by MmpL11 and MSMEG_0240 orthologues may represent a promising target for the development of new anti-mycobacterial compounds.

## Materials and Methods

### Bacterial strains and culture conditions

*Escherichia coli* TOP10 was used for the cloning steps and grown in Lysogeny Broth medium at 37°C supplemented with ampicillin (100 μg/ml), hygromycin (50 μg/ml) or kanamycin (25 μg/ml), when required. *Msm* mc^²^155 and the recombinant strains constructed in this study (Supplemental Table S1) were grown on Middlebrook 7H9 broth supplemented with glycerol (without OADC) or on 7H11 agar plates supplemented with glycerol and OADC enrichment, with hygromycin (50 μg/ml) and/or kanamycin (25 μg/ml), when required.

### BioID protocol with Msm cultures

Fifty ml of medium was inoculated at OD_600nm_=0.05 from 3-day saturated cultures of *Msm* and grown overnight (16 hours) in 7H9 medium with glycerol at 37°C with shaking. To avoid any depletion during the exponential phase of growth, biotin was added to a final concentration of 200 µM and the cultures were grown for a further 24 hours. After 40 hours of growth, the cultures were centrifuged at 4,000 x *g* for 10 min and then rinsed twice with Phosphate-Buffered Saline (PBS). Each pellet was resuspended in 900 µl of PBS per 400 mg of bacteria. Nine hundred µl of each culture were transferred to Lysing Matrix B tubes (MP Biomedicals) and the tubes were subjected to FastPrep (MP Biomedicals) treatment (6 cycles at maximum frequency for 1 min, 1 min of incubation on ice). One hundred µl of 10% n-DoDecyl-β-D-Maltoside diluted in PBS was added to the bacterial lysates and the mixture was further incubated for 2 hours at 4°C on a rotating wheel. After a quick spin, 700 µl of lysates were transferred to new Eppendorf tubes and centrifuged at 16,500 x *g* for 1 hr at 4°C. Dynabeads MyOne Streptavidin C1 (ThermoFisher Scientific) were rinsed three times with PBS, and 100 µl of beads were incubated with each clarified lysate at 4°C using a rotating wheel. After overnight incubation, the beads were washed three times with PBS containing 0.1% bovine serum albumin and twice with PBS (to reduce the bovine serum albumin signal during mass spectrometry analysis). Finally, the enriched biotinylated proteins were eluted with 100 µl of Laemmli buffer.

### Western-blot analysis

Twenty µL of total protein extract was resolved by SDS-PAGE on 12% acrylamide gels and then electrotransferred to a nitrocellulose membrane. The membrane was saturated with Tris-Buffered Saline (TBS), pH 7.5, containing 0.05% Tween 80 and 5% milk and probed overnight at 4 °C with anti-HA monoclonal antibody (Sigma-Aldrich, dilution 1:1,000) or with anti-Myc monoclonal antibody (Thermo Fisher Scientific, dilution 1:1,000) in TBS-Tween-3% milk. Finally, the membrane was incubated for 1 hr at room temperature with goat anti-mouse horseradish peroxidase (HRP)-conjugated secondary antibodies (Abcam, diluted 1:5,000) in TBS-Tween-3% milk. Alternatively, the biotinylated proteins were probed directly with HRP-conjugated streptavidin (BD Pharmingen, dilution 1:1,000) overnight. Blots were developed using the Amersham ECL Prime Western-Blotting Detection Reagent (GE Healthcare) and chemiluminescence was detected using the Amersham Imager 600 (GE Healthcare).

### Generation of Msm strains expressing APEX2 in the cytosol, cell wall, or as fusions to MmpL11

To express translational fusions of MmpL11 to APEX2 from an episomal plasmid, full-length MmpL11 and Nter MmpL11 were subcloned from pMV361_*mmpL11-birA* using primer pairs omlp1131/omlp1132 and omlp1131/omlp1133, respectively (Supplemental Table S2). The resulting PCR products and the vector pRibo-mpt63-APEX2m ^15^ were digested with NdeI and BamHI and ligated to yield pRibo-MmpL11-APEX2 and pRibo-Nter-APEX2. Both constructs were confirmed by Sanger sequencing. *Msm* was electroporated with pRibo-APEX2m, pRibo-mpt63-APEX2m ^14^, pRibo-MmpL11-APEX2, pRibo-Nter-APEX2 to yield strains Cyt-APEX2, Sec-APEX2, MmpL11-APEX2, and Nter-APEX2 (Supplemental Table S1). Cell stocks were made from individual clones obtained on selective agar (7H11 Middlebrook agar with 25 µg/mL kanamycin) and cultured in 7H9 Middlebrook with 25 µg/mL kanamycin.

### APEX2 mediated labelling and enrichment of biotinylated proteins

*Msm* strains Cyt-APEX2, Sec-APEX2, MmpL11-APEX2 and Nter-APEX2 were grown from stocks in 7H9 Middlebrook medium with 25 µg/mL kanamycin as previously described ^14^. Briefly, all strains were cultured in liquid medium to an optical density at 600 nm (OD_600_) of 0.8–1. Strains were then subcultured to OD_600_ 0.25 in 50 mL (Cyt-APEX2, Sec-APEX2, MmpL11-APEX2) or 200 mL (Nter-APEX2) with or without 2 mM theophylline inducer and cultured for an additional 2 generations (6 h). After cells were harvested by centrifugation at 2500 x*g* for 10 min, the pellets were resuspended at 10X concentration in 5 mL (all strains with culture volume 50 mL) or 20 mL (Nter-APEX2 culture volume 200 mL) in fresh medium containing 1 mM biotin-phenol (Iris Biotech) and incubated at 37°C for 30 min. Cell suspensions obtained after treatment with biotin-phenol were then treated with 1 mM H_2_O_2_ for 1 min. The reaction was quenched with an equal volume of 2X Quencher (20 mM sodium ascorbate, 20 mM sodium azide, 10 mM Trolox in PBS with 0.05% v/v Tween-80 [PBST80]). Cells were then washed sequentially with an equal volume of 1X quencher; equal volume of PBST80; 1.5X volume of 1X Quencher; and 1.5X volume of PBS.

Cell pellets obtained after the final wash were resuspended in PBS and lysed by bead-beating. Lysates were then clarified by centrifugation at 10,000 x *g* at 4°C and then treated with a final concentration of 1% (v/v) DDM at 4°C for 1 h to solubilise membrane fractions and extract membrane associated proteins. Biotinylated proteins were enriched by incubating lysates with neutravidin beads (Pierce PI29201) pre-equilibrated in PBS with gentle mixing at 22°C for 1 h, and then washing the beads three times with equal volumes of PBS. Enrichment and yield were confirmed by analysing a fraction of the beads by SDS-PAGE followed by streptavidin blot (Streptavidin IR-Dye 680 LT; LI-COR Odyssey detection) and silver stain (Pierce).

### Mass spectrometry proteomic analysis

Mass spectrometry proteomic analysis was performed as previously described ^36^. Raw data collected during nanoLC-MS/MS analyses were processed and converted to a *.mgf peak list format using Proteome Discoverer 2.5 (Thermo Fisher Scientific). MS/MS data were analysed using the Mascot search engine (version 2.4.0, Matrix Science, London, UK) installed on a local server. Searches were performed with a mass measurement tolerance of 10 ppm for precursor ions and 0.02 Da for fragment ions, against a composite target-decoy database constructed using a *Msm* mc²155 UniProt database (strain ATCC 700084/mc(2)155, taxo 246196, July 20172021, 12666 entries) fused with recombinant trypsin and a list of classical contaminants (118 entries). Cysteine carbamidomethylation, methionine oxidation, protein N-terminal acetylation and cysteine propionamidation were searched for as variable modifications. Up to one missed trypsin cleavage was allowed. Identification results were imported into Proline 2.0 software installed on a local server (http://proline.profiproteomics.fr) for validation ^37^. Peptide spectral matches greater than nine residues and ion scores >10 were retained. The false discovery rate was then optimised to less than 2% at the protein level using the Mascot Modified MudPIT score. Spectral counting analyses were performed using Proline.

For BioID analyses, we introduced a correction in order to compare the results as the total number of spectra can vary between independent biological replicates. For each condition, the number of spectra for a protein was normalised to Replicate 1 and was calculated as follows: Normalised number of spectra (Protein X) = Number of spectra (Protein X) in Replicate 1 + [Number of spectra (Protein X) in Replicate 2 * Total number of spectra in Replicate 1 / Total number of spectra in Replicate 2] + [Number of spectra (Protein X) in Replicate 3 * Total number of spectra in Replicate 1 / Total number of spectra in Replicate 3]. Since *birA* expression alone can lead to non-specific biotinylation of mycobacterial proteins ^16, 17^, the ratio was calculated as follows: Ratio (Protein X) = Normalised number of spectra with the protein of interest (MmpL11 or MSMEG_0240) in fusion with BirA / Normalised number of spectra with BirA alone. For calculation purposes, if the Normalised number of spectra with BirA alone for Protein X was zero, it was arbitrarily set to 0.5 (between 0 and 1 spectrum) in order to calculate a ratio. Only ratios greater than 10 were considered as meaningful, indicating that Protein X is enriched in this specific condition compared to the control (BirA alone). Since heterologous expression of the protein of interest alone could also potentially lead to non-specific biotinylation of proteins, the ratio of the protein of interest fused to BirA/BirA should be at least three times higher than the ratio of the protein of interest alone/BirA.

### Generation of SAINTexpress Analysis Portal

To facilitate accessible, high-confidence scoring of protein-protein interactions from affinity purification-mass spectrometry (AP-MS) data, we developed an interactive web-based analysis portal available at www.saintexpress.org. The front-end of the portal was built using the Angular framework, providing a user-friendly interface for data submission. Users can supply the standard input files (interaction, prey, and bait files) and select the appropriate quantification method matching their experimental design—supporting both spectral counts (SPC) and MS1-based intensity values (INT). Upon user-initiated execution, the front-end transmits the data to the back-end infrastructure built with Python and FastAPI, deployed behind an Nginx reverse proxy and hosted on Oracle Cloud Infrastructure (OCI). The back-end securely executes the analysis inside an isolated Docker container running the underlying SAINTexpress scoring algorithm. Once computation is complete, the resulting scored interaction data or error logs are returned to the user interface for review and download. To ensure computational transparency and reproducibility, the complete source code for the portal is publicly available on GitHub at https://github.com/dheerajramchandani/saintexpress-app. To protect user data privacy and manage server resources efficiently, all uploaded session files are removed immediately upon user-initiated session deletion, and a scheduled job automatically purges any remaining residual data from the server every 24 hours.

## Acknowledgments

This work was supported by an ANRS-MIE grant (decision 24579) to R.V.-C. and by Stony Brook University Renaissance School of Medicine funds to J.C.S.

## Abbreviations Used

BioID: proximity-dependent biotin identification
DDM: dodecyl maltoside
HRP: horseradish peroxidase
*Msm*: *Mycobacterium smegmatis*
*Mtb*: *Mycobacterium tuberculosis*
PPIs: protein-protein interactions

---

**Supplemental Figure S1. The model of interaction between the orthologues of MSMEG_0240 and the cytoplasmic domain of MmpL11 is conserved among mycobacteria.** AlphaFold3 prediction ^21^ of the interaction between the orthologue of MSMEG_0240 in (A) *M. tuberculosis*, (B) *M. leprae*, (C) *M. bovis* and (D) *M. marinum*, respectively, and the cytoplasmic domain of MmpL11.

**Supplemental Figure S2. APEX2 activity does not yield detectable biotinylation of MSMEG_0240.** Strains of *Msm* with MSMEG_0240 fused C-terminally to 1XFLAG and 3xHis epitopes at the native locus and expressing the indicated theophylline (theo)-inducible APEX2 constructs were subjected to the APEX2 labelling protocol with biotin-phenol. Biotinylated proteins were enriched by avidin enrichment. Biotinylated proteins were detected by avidin (*top*) and MSMEG_0240-1xFLAG-2xHis (asterisk; predicted MW: 27 kDa) was detected by anti-FLAG immunoblot (*bottom*) as previously reported by Ganapathy *et al*. ^23^. I, input; O, output.

**Supplemental Table S1. List of strains and plasmids used in this study.**

**Supplemental Table S2. List of primers used in this study.**

**Supplemental Table S3**. Mass spectrometry analysis of enriched biotinylated proteins from lysates of *Msm* mc²155 transformed with empty vector pMV361 or producing BirA, Nter, Nter-BirA, MmpL11 and MmpL11-BirA, when grown in 7H9 medium supplemented with glycerol (from three independent biological replicates).

**Supplemental Table S4.** Mass spectrometry analysis of enriched biotinylated proteins from lysates of *Msm* mc²155 producing BirA, MSMEG_0240, BirA-MSMEG_0240 and MSMEG_0240-BirA, when grown in 7H9 medium supplemented with glycerol (from three independent biological replicates).

**Supplemental Table S5**. Mass spectrometry analysis of enriched biotinylated proteins from lysates of *Msm* mc2155 encodeing Cyt-APEX2, Sec-APEX2, MmpL11-APEX2 or Nter-APEX2, when grown in 7H9 medium supplemented with glycerol (from three independent biological replicates).

**Supplemental Table S6**. SAINT analysis and annotation of results in Table S5 comparing biotinylated proteins enriched from *Msm* expressing APEX2 variants.

**Supplemental Table S7**. SAINT analysis and annotation of results in Tables S3 comparing biotinylated proteins enriched from *Msm* expressing MmpL11-BirA or Nter-BirA.

**Supplemental Table S8**. SAINT analysis and annotation of results in Tables S4 comparing biotinylated proteins enriched from *Msm* expressing BirA-MSMEG_0240 or MSMEG_0240-BirA.

