## Supplementary material for "The cytoplasmic C-terminal domain of the MmpL11 lipid transporter is required for interaction with its co-cistronic partner MSMEG_0240 in *Mycobacterium smegmatis*": FigS1 and S2, Tables S1 and S2

**Supplemental Information**

**Supplemental Table S1.** List of strains and plasmids used in this study.

| **Strains and plasmids** | **Genotype or description** | **Source or reference** |
| --- | --- | --- |
| *E. coli* TOP10 | F^-^ *mcr*A Δ(*mrr*-*hsd*RMS-*mcr*BC) φ80*lac*Z ΔM15 Δ*lac*X74 *rec*A1 *ara*D139 Δ(*ara* *leu*)7697 *gal*U *gal*K *rps*L (StrR) *end*A1 *nup*G, used for cloning | Invitrogen |
| *M. smegmatis* mc²155 |  | ATCC 700084 |
| Δ*MSMEG_0241* (Δ*mmpL11*) | *M. smegmatis* mc²155 Δ*MSMEG_0241*::*hygR* | ^1^ |
| pcDNA3.1 mycBioID | Plasmid used to fuse any protein of interest to the C-terminus of BirA(R118G) with a Myc tag | Addgene #35700, ^2^ |
| pcDNA3.1 MCS-BirA(R118G)-HA | Plasmid used to fuse any protein of interest to the N-terminus of BirA(R118G) with a HA tag | Addgene #36047, ^2^ |
| pMV361 | Monocopy *E. coli* - mycobacteria shuttle vector, *hsp60* promoter, integrative at *attB* site | ^3^ |
| pcDNA_*birA*-*Nter* | Plasmid allowing the production of truncated MSMEG_0241 (Nter) fused to the C-terminus of BirA(R118G) with a Myc tag in *E. coli* | This work |
| pcDNA_*birA*-*mmpL11* | Plasmid allowing the production of MSMEG_0241 fused to the C-terminus of BirA(R118G) with a Myc tag in *E. coli* | This work |
| pcDNA_*birA*-*Cter* | Plasmid allowing the production of truncated MSMEG_0241 (Cter) fused to the C-terminus of BirA(R118G) with a Myc tag in *E. coli* | This work |
| pcDNA_*Nter*-*birA* | Plasmid allowing the production of truncated MSMEG_0241 (Nter) fused to the N-terminus of BirA(R118G) with a HA tag in *E. coli* | This work |
| pcDNA_*mmpL11*-*birA* | Plasmid allowing the production of MSMEG_0241 fused to the N-terminus of BirA(R118G) with a HA tag in *E. coli* | This work |
| pcDNA_*birA*-*MS0240* | Plasmid allowing the production of MSMEG_0240 fused to the C-terminus of BirA(R118G) with a Myc tag in *E. coli* | This work |
| pcDNA_*MS0240*-*birA* | Plasmid allowing the production of MSMEG_0240 fused to the N-terminus of BirA(R118G) with a HA tag in *E. coli* | This work |
| pMV361_*birA* | Integrative plasmid allowing the production of *E. coli* biotin ligase BirA(R118G) with a HA tag in mycobacteria | ^4^ |
| pMV361_*Nter* | Integrative plasmid allowing the production of truncated MSMEG_0241 in mycobacteria | This work |
| pMV361_*mmpL11* | Integrative lasmid allowing the production of MSMEG_0241 in mycobacteria | This work |
| pMV361_*birA*-*Nter* | Integrative plasmid allowing the production of truncated MSMEG_0241 (Nter) fused to the C-terminus of BirA(R118G) with a Myc tag in mycobacteria | This work |
| pMV361_*birA*-*mmpL11* | Integrative plasmid allowing the production of MSMEG_0241 fused to the C-terminus of BirA(R118G) with a Myc tag in mycobacteria | This work |
| pMV361_*birA*-*Cter* | Integrative plasmid allowing the production of truncated MSMEG_0241 (Cter) fused to the C-terminus of BirA(R118G) with a Myc tag in mycobacteria | This work |
| pMV361_*Nter*-*birA* | Integrative plasmid allowing the production of truncated MSMEG_0241 (Nter) fused to the N-terminus of BirA(R118G) with a HA tag in mycobacteria | This work |
| pMV361_*mmpL11*-*birA* | Integrative plasmid allowing the production of MSMEG_0241 fused to the N-terminus of BirA(R118G) with a HA tag in mycobacteria | This work |
| pMV361_*MS0240* | Integrative plasmid allowing the production of MSMEG_0240 in mycobacteria | This work |
| pMV361_*birA*-*MS0240* | Integrative plasmid allowing the production of MSMEG_0240 fused to the C-terminus of BirA(R118G) with a Myc tag in mycobacteria | This work |
| pMV361_*MS0240*-*birA* | Integrative plasmid allowing the production of MSMEG_0240 fused to the N-terminus of BirA(R118G) with a HA tag in mycobacteria | This work |
| pRibo-APEX2m | Episomal plasmid used for the theophylline-inducible expression of Cyt-APEX2 in mycobacteria | Addgene #176842, ^5^ |
| pRibo-Sec-APEX2m | Episomal plasmid used for the theophylline-inducible expression of Sec-APEX2 in mycobacteria | Addgene #176844, ^5^ |
| pRibo-MmpL11-APEX2 | Episomal plasmid used for the theophylline-inducible expression of MmpL11-APEX2 fusion in mycobacteria | This work |
| pRibo-Nter-APEX2 | Episomal plasmid used for the theophylline-inducible expression of the N-terminus of MmpL11 fused to APEX2 in mycobacteria | This work |

**Supplemental Table S2.** List of primers used in this study.

| **Primers** | **5' to 3' sequence** |
| --- | --- |
| pcDNA_*birA-mmpL11*_dir | TATATA**GAATTC** cgcttgagcagcactttgcgca (**EcoRI**) |
| pcDNA_*birA-Nter*_rev | TATA**AAGCTT**TCA caccagcacgagccgcacc (**HindIII**) |
| pcDNA_*birA-mmpL11*_rev | tata**aagctt** tcacttcgcctcctccagcattgc (**HindIII**) |
| pcDNA_*birA-Cter*_dir | tata**gaattc** ccggcgctcatggcgatgttc (**EcoRI**) |
| pcDNA_*mmpL11-birA*_dir | TATA**GCTAGC**CAATTG tgatgcgcttgagcagcactttgc (**NheI** and MunI) |
| pcDNA_*Nter-birA*_rev | TATATA**GAATTC** caccagcacgagccgcacc (**EcoRI**) |
| pcDNA_*mmpL11-birA*_rev | TATATA**GAATTC** cttcgcctcctccagcattgctacc (**EcoRI**) |
| pcDNA_*birA*-*MS0240*_dir | TATATA**GCGGCCGC** tggcccccgacgttcggg (**NotI**) |
| pcDNA_*birA*-*MS0240*_rev | TATA**AAGCTT** tcacttgccgctgcgctgc (**HindIII**) |
| pcDNA_*MS0240-birA*_dir | TATA**GCTAGC**CAATTG tggcccccgacgttcggg (**NheI** and MunI) |
| pcDNA_*MS0240-birA*_rev | TATATA**TCCGGA** cttgccgctgcgctgcgc (**Kpn2I**) |
| pMV361_*mmpL11*_dir | TATA**CAATTG** tgatgcgcttgagcagcactttgc (**MunI**) |
| pMV361_*Nter*_rev | TA**GTTAAC** caccagcacgagccgcacc (**HpaI**) |
| pMV361_*mmpL11*_rev | TA**GTTAAC** tcacttcgcctcctccagcattg (**HpaI**) |
| pMV361_*MS0240*_dir | TATA**CAATTG** tggcccccgacgttcggg (**MunI**) |
| pMV361_*MS0240*_rev | TA**GTTAAC** tcacttgccgctgcgctgc (**HpaI**) |
| pMV361_*birA*-xxx_dir | CCATGGAACAAAAACTCATCTCAGAAGAGG |
| Up_*MS0240*_dir | TATATA**actagt** ggttgtcggtggccgtcgac (**SpeI**) |
| Up_*MS0240*_rev | TATATA**gatatc** gacatgtctttgcgacgcgcgtc (**EcoRV**) |
| Down_*MS0240*_dir | TATATA**tctaga** ggttgtcggaccggttcgcg (**XbaI**) |
| Down_*MS0240*_rev | TATATA**aggcct** cgaaccggcgcagcgagac (**StuI**) |
| Seq_*MS0240*_dir (1) | cacccgcctgtcgacgcag |
| Seq_*hygro*_rev (2) | CAGGACCTGCAGGCATGCAAGC |
| Seq_*hygro*_dir (3) | GCCCCCGGCGCCTGACG |
| Seq_*MS0240*_rev (4) | cgggtgcggtcggtgacc |
| omlp1131 (mmpL11 fwd) | GCAACAAGATGCATATGATGCGCTTGAGCAGCAC |
| omlp1132 (mmpL11 rev) | AGCTCTTGCCGGATCCGAATTCCTTCGCCTCCTCCAGCATTG |
| omlp1133 (Nter mmpL11 rev) | AGCTCTTGCCGGATCCGAATTCCACCAGCACGAGCCGC |
| omlp1215 | gtcgctgctgctgcggatgcagcggttgtcggaccggttcgcgaagatcgccgcgcagcgcagcggcaag  GGTTTGTCTGGTCAACCACCGCGGTCTCAGTGGTGTACGGTACAAACCtgatccggcaggcggattcggttctggccgagcggatggcttgactcacacccacgccacgggtactgat |
| omlp1216 | GTATCTGGAGGGCAGTGTGAG |
| omlp1217 | GTTCGGTCAGTCACGAGTAG |

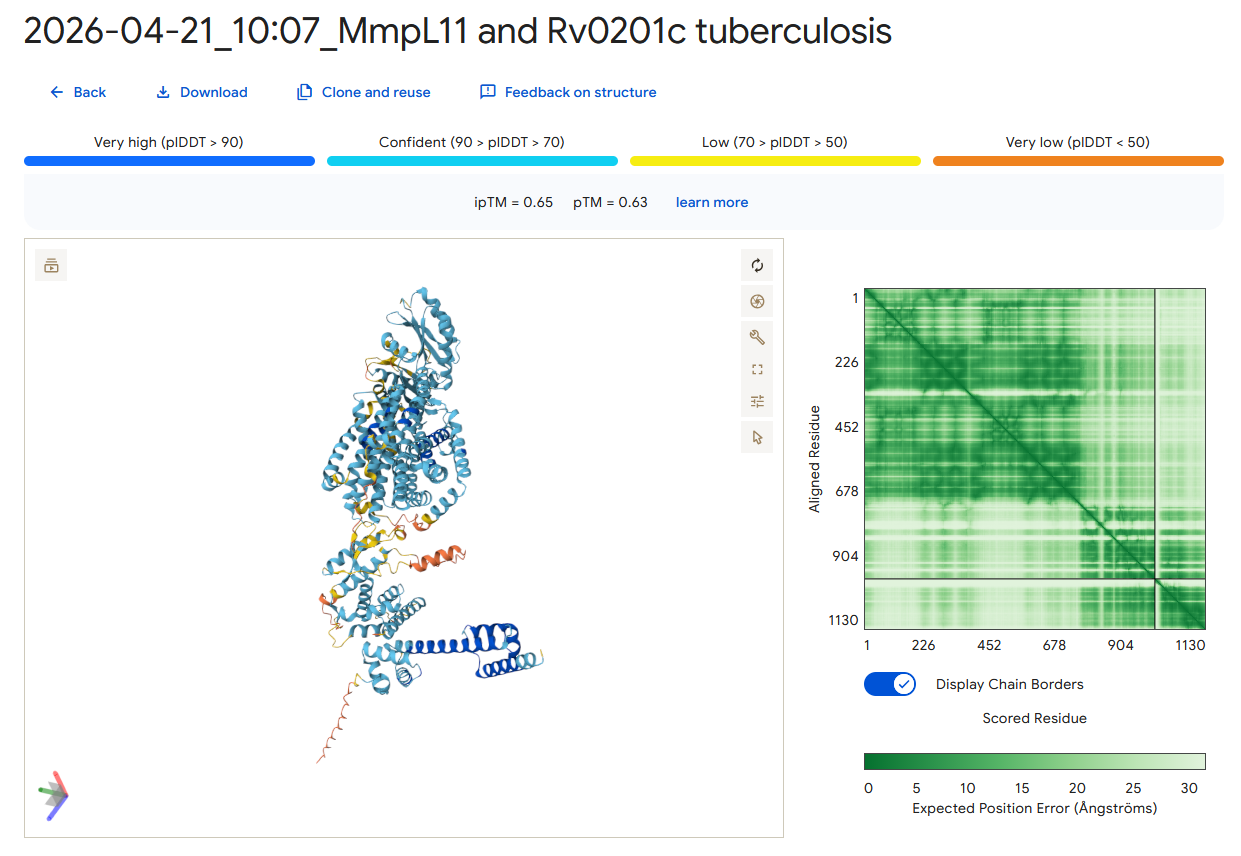
**A**

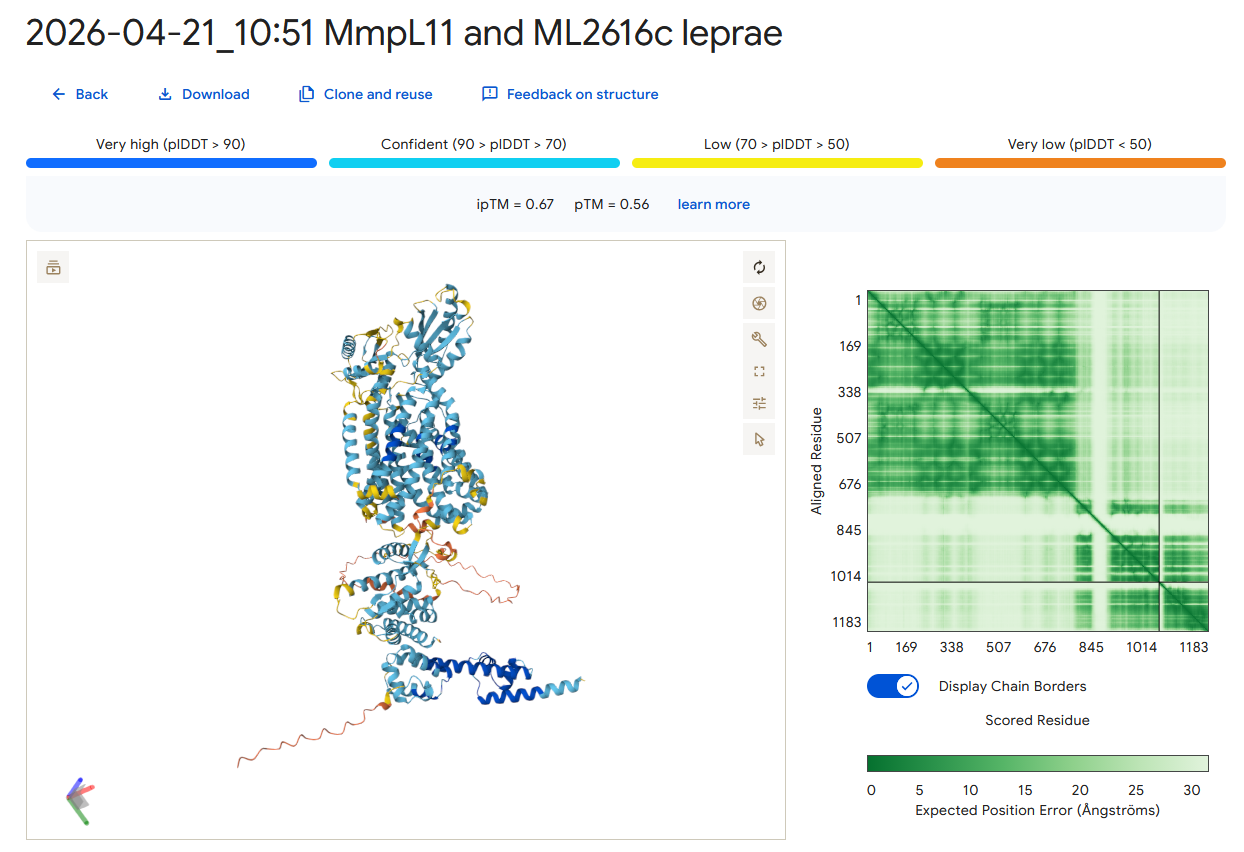
**B**

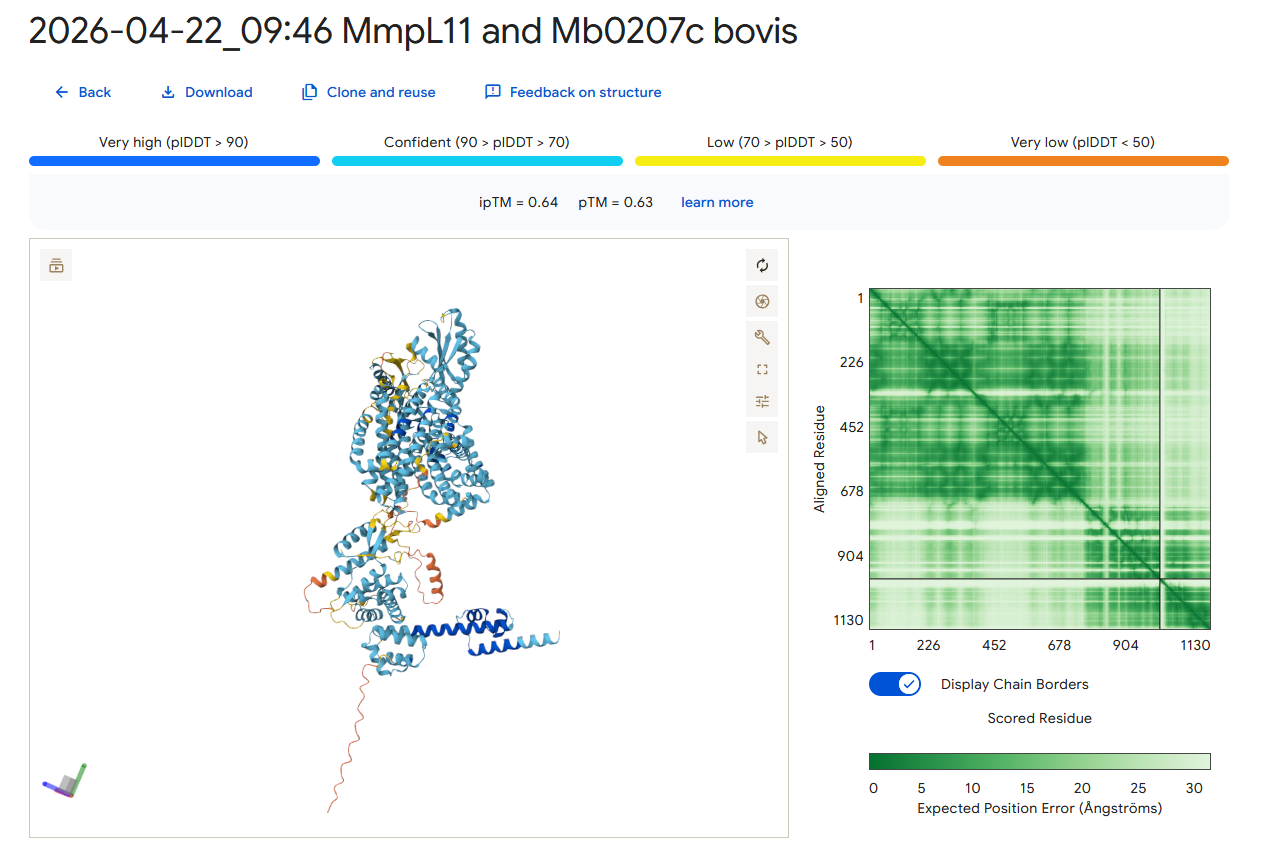
**C**

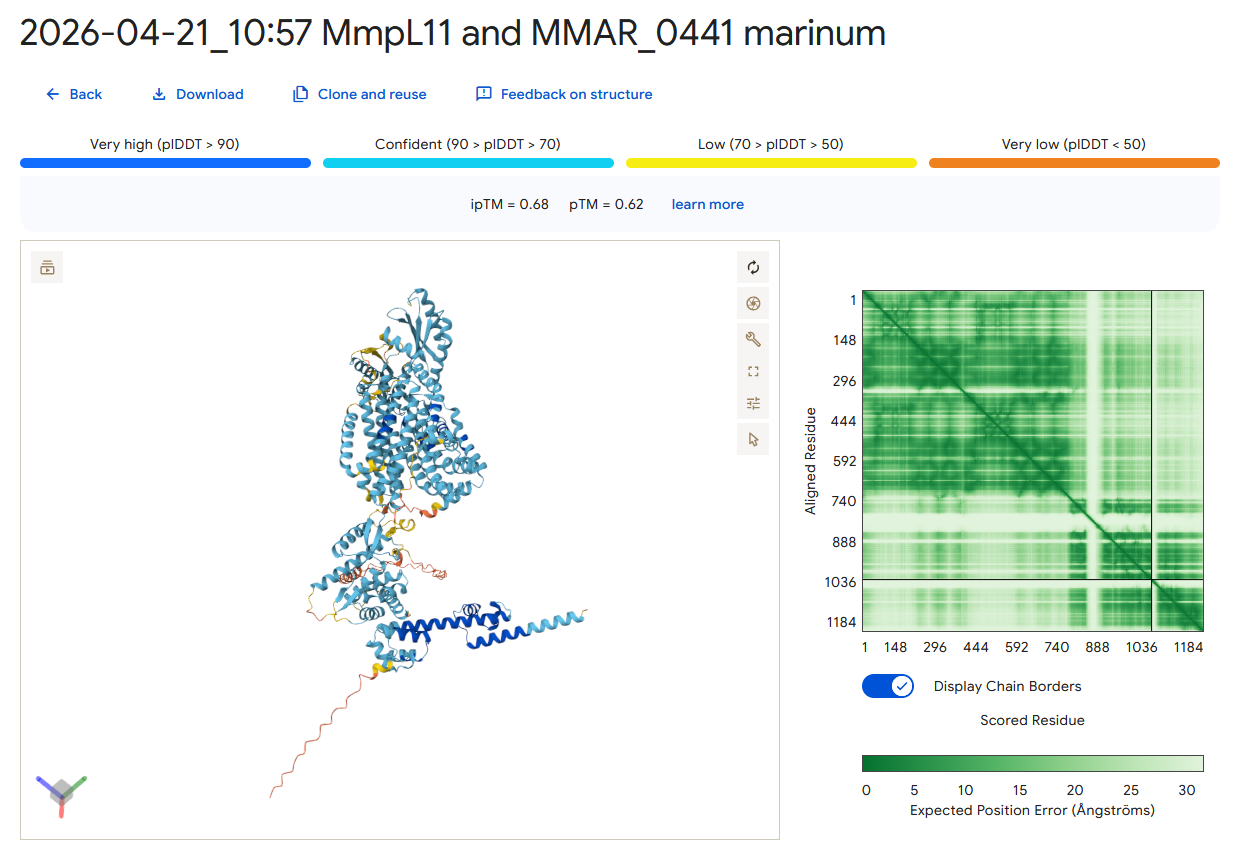
**D**

**Supplemental Figure S1. The model of interaction between the orthologues of MSMEG_0240 and the cytoplasmic domain of MmpL11 is conserved among mycobacteria.** AlphaFold3 prediction^6^ of the interaction between the orthologue of MSMEG_0240 in **(A)** *M. tuberculosis* (Rv0201c), **(B)** *M. leprae* (ML2616c), **(C)** *M. bovis* (Mb0207c) and **(D)** *M. marinum* (MMAR_0441), and the cytoplasmic domain of MmpL11**.**

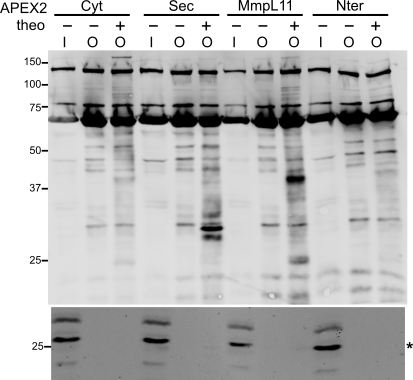

**Supplemental Figure S2. APEX2 activity does not yield detectable labelling of MSMEG_0240.** Strains of *Msm* with MSMEG_0240 fused C-terminally to 1XFLAG and 3xHis epitopes at the native locus and expressing the indicated theophylline (theo)-inducible APEX2 constructs were subjected to the APEX2 labeling protocol with biotin-phenol. Biotinylated proteins were enriched by avidin enrichment. Biotinylated proteins were detected by avidin (*top*) and MSMEG_0240-1xFLAG-2xHis (asterisk; predicted MW: 27 kDa) was detected by anti-FLAG immunoblot (*bottom*) as previously reported by Ganapathy *et al*. ^6^. I, input; O, output.

**Supplemental Methods**

*Genetic constructions*

MmpL3, MmpL11 and MmpL13 are closely related. The *Msm* homologue of MmpL11, MSMEG_0241, was assigned not only by sequence identity, but also gene synteny (data not shown). Two types of constructs were used for MmpL11: a truncated version, including the 12 TMD and lacking the C-terminal cytoplasmic domain (hereafter referred to as Nter), or the full-length MmpL11 from *Msm* mc²155. Therefore, the *mmpl11* and *MSMEG_0240* genes were amplified by PCR from *Msm* mc²155 genomic DNA using primers pMV361_*mmpL11*_dir and pMV361_*Nter*_rev for the truncated version of *mmpL11*, or pMV361_*mmpL11*_dir and pMV361_*mmpL11*_rev for the full-length *mmpL11*, or pMV361_*MS0240*_dir and pMV361_*MS0240*_rev for *MSMEG_0240* (Supplemental Table S2). The corresponding amplicons were digested with MunI and HpaI, and inserted into pMV361 digested with the same enzymes, to give pMV361_*Nter*, pMV361_*mmpL11* and pMV361_*MS0240*. The corresponding proteins are shown in Figure 1A (constructs 1 and 2) and Figure 2A (construct 1).

To generate the vectors expressing the fusion between *Nter* or *mmpL11* or *Cter* and *birA*, the genes were amplified by PCR from *Msm* mc²155 genomic DNA using the primers pcDNA_*birA-mmpL11*_dir and pcDNA_*birA-Nter*_rev, or pcDNA_*birA-mmpL11*_dir and pcDNA_*birA-mmpL11*_rev, or pcDNA_*birA-Cter*_dir and pcDNA_*birA-mmpL11*_rev or pcDNA_*mmpL11-birA*_dir and pcDNA_*Nter-birA*_rev or pcDNA_*mmpL11-birA*_dir and pcDNA_*mmpL11-birA*_rev (Supplemental Table S2). For the *birA-Nter*, *birA-mmpL11* and *birA-Cter* fusions, the *Nter*, *mmpL11* and *Cter* amplicons were digested with HindIII and EcoRI and inserted into pcDNA3.1 mycBioID (Addgene, ^2^) digested with the same enzymes to give pcDNA_*birA-Nter*, pcDNA_*birA-mmpL11* and pcDNA_*birA-Cter*, respectively. The *birA-Nter*, *birA-mmpL11* and *birA-Cter* fragments were amplified from pcDNA_*birA-Nter*, pcDNA_*birA-mmpL11* and pcDNA_*birA-Cter*, using primers pMV361_*birA-xxx*_dir and pcDNA_*birA-Nter*_rev, or pMV361_*birA-xxx*_dir and pcDNA_*birA-mmpL11*_rev (Supplemental Table S2), digested with HindIII and inserted into pMV361, previously digested with HindIII and PvuII, to give pMV361_*birA-Nter*, pMV361_*birA-mmpL11* and pMV361_*birA-Cter*. The corresponding fusion proteins are shown in Figure 1A (constructs 3, 4 and 5). For the *Nter-birA* and *mmpL11-birA* fusions, the *Nter* and *mmpL11* amplicons were digested with EcoRI and NheI and inserted into pcDNA3.1 MCS-BirA(R118G)-HA (Addgene, ^2^) digested with the same enzymes to give pcDNA_*Nter-birA* and pcDNA_*mmpL11-birA*, respectively. The *Nter-birA* and *mmpL11-birA* fragments were obtained after digestion of pcDNA_*Nter-birA* and pcDNA_*mmpL11-birA* with MunI and PmeI and inserted into pMV361, digested with MunI and HpaI, to give pMV361_*Nter-birA* and pMV361_*mmpL11-birA*. The corresponding fusion proteins are shown in Figure 1A (constructs 6 and 7).

To generate the vectors expressing the fusion between *MSMEG_0240* and *birA*, the gene was amplified by PCR from *Msm* mc²155 genomic DNA using primers pcDNA_*birA-MS0240*_dir and pcDNA_*birA-MS0240*_rev, or pcDNA_*birA-MS0240*_dir and pcDNA_*birA-MS0240*_rev (Supplemental Table S2). For the *birA-MSMEG_0240* fusion, the *MS0240* amplicon was digested with HindIII and NotI and inserted into pcDNA3.1 mycBioID (Addgene, ^2^) digested with the same enzymes to give pcDNA_*birA-MS0240*. The *birA-MS0240* fragment was amplified from pcDNA_*birA-MS0240*, using primers pMV361_*birA-xxx*_dir and pcDNA_*birA-MS0240*_rev (Supplemental Table S2), digested with HindIII and inserted into pMV361, previously digested with HindIII and PvuII, to give pMV361_*birA-MS0240*. The corresponding fusion protein is shown in Figure 2A (construct 2). For the *MSMEG_0240-birA* fusion, the *MS0240* amplicon was digested with Kpn2I and NheI and inserted into pcDNA3.1 MCS-BirA(R118G)-HA (Addgene, ^2^), which was digested with the same enzymes to give pcDNA_*MS0240-birA*. The *MS0240-birA* fragment was obtained after digestion of pcDNA_ *MS0240-birA* with MunI and PmeI and inserted into pMV361, digested with MunI and HpaI, to give pMV361_*MS0240-birA*. The corresponding fusion protein is shown in Figure 2A (construct 3). All the constructions were verified by DNA sequencing before being into electrocompetent cells of *Msm* mc²155.

To generate constructs for expression of APEX2 fusions of either Nter or full length MmpL11, PCR fragments were amplified from the plasmid pMV361_*birA*-*mmpL11* using oligos omlp1131 & omlp1132 and omlp1131 & omlp1133, respectively. PCR fragments obtained as above and the plasmid pRibo-mpt63-APEX2 were digested with Nde1 and BamH1. Ligation of the respective digested PCR fragments with the digested APEX2 plasmid was performed to obtain the following plasmids: pRibo-MmpL11-APEX2 and pRibo-Nter-APEX2.

*Construction of Msm with MSMEG_0240-1xFLAG-2xHis by ORBIT*

*Msm* with the MSMEG_0240 coding region C-terminally translational fused to 1xFLAG-3xHis was created using ORBIT as reported by Murphy *et al*. ^7^. Briefly, *Msm* mc^2^155 harboring the plasmid pKM461 (Addgene #108320) was grown to OD_600_ 0.5 and treated with 500 ng/mL anhydrotetracycline for an additional doubling time (~3 h). Electrocompetent cells prepared from this culture were electroporated with pKM491-FLAG-His (Addgene#109282) and the attP-containing targeting oligonucleotide omlp1215 (Table S1). Clones were selected on Middlebrook 7H10/glycerol agar (“7H10”) with 50 µg/mL hygromycin. Ten clones were selected and screened for loss of pKM461 by selecting on 7H10 containing 3% sucrose. Colonies were isolated and confirmed for growth on 7H10 with and without 25 µg/mL kanamycin. Insertion of the FLAG-His tag at the MSMEG_0240 C-terminus was verified by PCR of genomic DNA and sequencing of the PCR product using primers omlp1216 and omlp1217 (Table S1).
